# clinTALL: machine learning-driven multimodal subtype classification and treatment outcome prediction in pediatric T-ALL

**DOI:** 10.64898/2026.01.28.701965

**Authors:** Lukas Stoiber, Željko Antić, Stefano Rebellato, Grazia Fazio, Annika Rademacher, Lennart Lenk, Franco Locatelli, Adriana Balduzzi, Gunnar Cario, Carmelo Rizzari, Giovanni Cazzaniga, Jiangyan Yu, Anke Katharina Bergmann

## Abstract

**Background:** Childhood T-lineage acute lymphoblastic leukemia (T-ALL) is an aggressive hematologic malignancy with poor prognosis. Differently from B-cell precursor ALL, T-ALL lacks effective risk stratification strategies. A recent study has integrated whole genome and whole transcriptome data to define over 15 distinct molecular subtypes with prognostic significance. However, clinical translation of this knowledge remains challenging due to the complexity of interpreting high-dimensional multi-omics-based data.

**Methods:** Here, we present clinTALL, a deep learning based multi-task pipeline for pediatric T-ALL subtype classification and treatment outcome estimation. The model integrates multimodal input data and uses a neural network architecture to generate a shared latent embedding for jointly learned multi-task prediction. The competing risk-based model was used to predict event-specific outcomes. The model was trained on a publicly available multimodal dataset comprising clinical, genomic and transcriptomic features of 1309 pediatric T-ALL samples.

**Results:** We observed that the transcriptomic-only model achieved superior single modality results, with 92.2% accuracy for subtype prediction and a 65.9% concordance index (C-index) for event-free survival (EFS) in a cross-validation setup. Integrating all data modalities maintained high subtype classification accuracy (91.7%) and improved the overall concordance index for EFS estimation to 67.5%. The competing risk-based model enables accurate predictions of induction failure (C-index = 96.0%) and second malignant neoplasm (C-index = 62.1%). We validated molecular subtype predictions on an internal dataset of 120 pediatric T-ALL samples and obtained an accuracy of 81.8%. To facilitate the broad application of multi-omics based subtype prediction and treatment outcome inference, we provide clinTall as a Docker based application, allowing for user friendly access to the tool. The full source code of clinTALL is available on GitHub (https://github.com/UKWgenommedizin/clinTALL).

**Conclusion:** Together, our machine learning-based framework allows for automated, accurate sub-type classification and treatment outcome inference using multimodal input data, advancing precision risk stratification for pediatric T-ALL.

## Background

T-lineage acute lymphoblastic leukemia (T-ALL) is an aggressive hematologic malignancy with poor treatment outcome compared to B-lineage ALL [1, 2]. In contrast to the extensively characterized B-ALL, which has well defined genetic biomarkers that guide risk stratification and therapy, the molecular heterogeneity of T-ALL remains poorly characterised [3]. At present, the World Health Organization (WHO) and International Consensus Classification (ICC) recognize only early T-cell precursors ALL (ETP-ALL) as a distinct subtype [4], yet multi-omics studies have now substantially refined our understanding of T-ALL heterogeneity by defining up to 17 genomic subtypes with growing evidence of prognostic relevance [5–8].

Translating high dimensional multi-omics discoveries into clinical practice remains challenging. However, recent advances in machine learning (ML) have begun to bridge this gap in pediatric ALL. For example, several ML-based subtype classifiers have been developed that integrate one or multiple layers of genomic, transcriptomic or epigenomic features to define ALL molecular subtypes with increasing accuracy [9–13]. In addition, multi-omics data have been used for cause-specific survival modeling to identify patients with poor outcomes [8, 14, 15]. Although such cause-specific approaches can determine high risk groups, they do not explicitly predict which clinical endpoint is most likely to occur [16]. The endpoint specific prediction is critical in T-ALL, which contains distinct treatment outcome events, such as induction failure, relapse, toxic death and others, reflecting complex interaction between different biological mechanisms and insufficiently refined treatment options in driving these unfavorable events [17]. Accordingly, competing-risks methods, rather than single-event survival models, are required to enable accurate and event-specific outcome prediction.

Here we present clinTALL, a ML-based framework that integrates clinical and multi-omics data to enable both subtype classification and treatment outcome prediction in pediatric T-ALL. Multi-modal tabular data from 1309 pediatric T-ALL samples from a recent publication [8] were used to train a TabM-derived architecture for downstream multi-task learning [18], including genetic subtype identification and competing-risks modeling. Our framework achieves high accuracy of global subtype classification (91.7%), outperforming available tools, and demonstrates strong prediction accuracy in independent external cohorts (81.8%). The competing risk-based modeling enables event-specific risk prediction, and accurately identifies patients with increased risk for induction failure or second malignant neoplasm. Notably, it is able to determine patients with increased likelihood of relapse, which showed significantly reduced 5-years event free survival (EFS) probability (75.5% vs 85.7%, p = 7.03 × 10^−5^). By making clinTALL publicly available, we aim to facilitate diagnostic standardization, support translational efforts, and accelerate the incorporation of molecular sub-type information into pediatric T-ALL biology and patient management.

## Methods

### Datasets

#### Training cohort

The training cohort was derived from the dataset published by Pölönen et al. [8] (Children’s Oncology Group (COG) AALL0434 trial). The data were accessed via Synapse (ID: syn54032669; https://doi.org/10.7303/syn54032669). Whole-genome sequencing data and clinical annotations were extracted from the supplementary tables of the original publication.

#### Validation cohort I (AIEOP-BFM ALL 2017)

The first validation cohort comprised 120 diagnostic samples from patients with childhood T-ALL enrolled in the Italian AIEOP-BFM ALL 2017 (EudraCT Number: 2016-001935-12; NCT 03643276) treatment study. Whole-transcriptome sequencing was performed according to the manufacturer’s guidelines. Raw sequencing reads were quality-tested with FastQC [19] and aligned against the GRCh38/hg38 reference human genome with STAR aligner (v2.7.10b) [20]. The gene level count matrix was generated using featureCounts (v2.0.0) [21] for downstream clinTALL based prediction. GRCh38 Ensembl Release 108 annotation was used as reference.

#### Validation cohort II (NOPHO ALL2008)

The second validation cohort was derived from the study by Hackenhaar et al. [22] and included 108 diagnostic T-ALL samples from the Nordic Society of Paediatric Haematology and Oncology (NOPHO) ALL2008 trial. Gene expression data were retrieved from the Gene Expression Omnibus (GEO) under accession number GSE272023.

### Data Preprocessing

#### Cohort and data modalities

Multi-omics data integration was performed using a cohort of 1309 T-ALL samples [8]. Four data modalities were included: gene expression, copy number variation (CNV), DNA variants, and clinical data.

#### Sample filtering and label harmonization

Samples lacking outcome labels or survival information were excluded from all analyses. Two samples originally annotated as “induction death” were reassigned to the “death, not otherwise specified (NOS)” category. This reassignment was defined *a priori* to ensure numerical stability during model training and downstream analyses.

#### Feature construction and preprocessing

##### Gene expression

Gene expression features consisted of transcript-level expression values. Gene expression features were subjected to supervised feature selection to constrain dimensionality and reduce downstream model complexity. A Random Forest classifier (RandomForestClassifier, scikit-learn; 100 trees, maximum depth = 10; [23]) was trained to predict disease subtypes. Feature importance was quantified using SHAP (SHapley Additive exPlanations; [24]) with the TreeExplainer framework. Class-specific mean absolute SHAP values were used to rank features.

The top 100 genes per subtype were selected, yielding 474 unique genes across all subtypes (Supplementary Table S1). To mitigate information loss from aggressive filtering and facilitate learning of nonlinear feature interactions, 526 highly variable genes were reintroduced. This combined gene set was used for both gene expression–only and multimodal models.

##### Copy number variation

CNV features were derived from Supplementary Table 19 of the Pölönen et al. study [8] by aggregating CNV events per megabase. Features were stratified by CNV type (e.g., deletion, loss, gain) and size category (e.g., focal, small, broad). Low-information features were removed by excluding features with low variance (*<* 0.1) or fewer than 10 unique values, yielding a total of 458 CNV features.

##### DNA variants

DNA variant features were obtained from the Synapse R dataset (Data 1309Samples.RData, https://doi.org/10.7303/syn54032669) and augmented with variants from Supplementary Table 11 of Pölönen et al. [8], resulting in 608 variant features.

##### Clinical data

Four post-induction clinical variables (“Day 29 morphology”, “Treatment Arm”, “Standard Induction”, and “HSCT”) were excluded, resulting in a total of 42 encoded clinical features.

#### Model construction

Unimodal models were trained separately for gene expression, DNA variants, CNVs, and clinical data using the corresponding feature sets described above. The clinical-only model used all retained clinical variables, the variant-only model used all available variant features, and the CNV-only model used all filtered CNV features.

For multimodal integration, gene expression, DNA variant, and clinical features were combined. CNV features were excluded from the multimodal model due to consistently poor performance observed in unimodal CNV models, indicating limited predictive utility in this setting. For a detailed overview of features per model see supplementary tables S2 and S3.

#### Encoding

Categorical variables (e.g., CNS status) were one-hot encoded. Continuous variables (e.g., white blood cell count) were not standard-scaled; instead, they were processed using Piecewise Linear Embeddings (PLE; [25]) which represent scalar features through learnable piecewise linear transformations, improving numerical stability and expressiveness.

#### Deep Learning Architecture

We employed a TabM-derived architecture ([18]), a deep learning framework optimized for tabular data that utilizes a mini-ensemble technique within network layers. The architecture consisted of three components:

#### Embeddings

Numerical features were transformed using PLE with learnable quantiles, enabling modeling of nonlinear relationships without explicit pre-normalization.

#### Backbone

The core network used the TabM structure, parameterized by number of network layers (*n*_block_) and layer dimension (*d*_block_, width of layers).

#### Multi-task competing risk head

The network was designed for multi-task learning, simultaneously predicting:

- **Competing-risk survival**, modeled using a cause-specific Cox proportional hazards framework, producing *K* log-hazard outputs corresponding to distinct event types (e.g., relapse).
- **Subtype classification**, modeled as a multiclass classification task.

#### Training and Stratification Strategy

Model training employed a custom relaxed stratified 5-fold cross-validation scheme. To address class imbalance and correlations between subtypes and survival outcomes, a hierarchical stratification strategy was applied. Samples were initially stratified by a composite *Subtype EventStatus* label; rare categories were pooled into an “OTHER” group when necessary.

Data were split into outer 5-fold cross-validation sets, with each training fold further divided into training and validation subsets (80%/20%). Training used the AdamW optimizer [26] with full-batch gradient descent. The global loss was defined as:

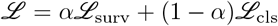

where ℒ_surv_ denotes the sum of cause-specific Cox partial likelihood losses (Breslow method for ties) and ℒ_cls_ denotes the cross-entropy loss. Inverse class-frequency weights were (optionally) applied to both loss components. Early stopping was based on the sum of validation accuracy and concordance index (C-index) with a patience of 50 epochs.

#### Hyperparameter Optimization

Hyperparameter tuning was conducted using Optuna [27] with 75 trials per dataset configuration. Optimized parameters included learning rate (10^−5^ – 10^−2^), weight decay (10^−6^ – 10^−3^),architecture dimensions (block sizes: 128, 256, 512; embedding sizes: 8, 12, 16), number of PLE bins (16, 32, 64), PLE function usage (true/-false), loss weighting parameter *α* (0.1 – 0.9), and inverse class-frequency weighting (true/false). A list of final model parameters can be found in supplementary table S4.

The objective function was a weighted average of threefold cross-validation C-index and classification accuracy, with equal weight assigned to each metric. For an overview of final model parameters after tuning see supplementary table S4.

#### Statistical Analysis and Implementation

Models were implemented in PyTorch [28] using the rtdl-num-embeddings and TabM library ([18, 25]). Survival metrics were computed using scikit-survival ([29]). Experiments were run on a CUDA-enabled GPU. Performance was reported as mean C-index and accuracy across the five cross-validation folds. Analyses were conducted in Python using SciPy [30], lifelines [31], scikit-survival [29], matplotlib [32], and seaborn [33].

#### Competing-Risk Cumulative Incidence Analysis

Cause-specific cumulative incidence functions were estimated for five mutually exclusive event types: relapse, toxic death, induction failure, second malignant neoplasm, and death NOS using scikit-survival’s `cumulative_incidence_competing_risks` function. Patients were stratified into high- and low-risk groups based on predicted cause-specific risk scores, using percentile-based thresholds corresponding to observed event incidences.

#### Event-Free Survival Analysis

Event-free survival (EFS) was analyzed to assess the prognostic relevance of predicted relapse risk. Patients were stratified into high- and low-risk groups using the upper decile of predicted relapse risk. Survival time was administratively censored at five years. Kaplan–Meier curves were estimated using lifelines `KaplanMeierFitter`, and differences between groups were evaluated using the log-rank test.

#### Benchmarking clinTALL and TALLSorts

To establish baseline performance, we evaluated the TALLSorts classifier [9] using repeated cross-validation. A custom Python wrapper enabled automated training and evaluation via the TALLSorts command-line interface. Only SHAP-preselected genes were used to limit runtime.

Five-fold cross-validation was repeated for 10 independent iterations with distinct random seeds (50 total evaluations). Models were trained using default TALLSorts parameters and evaluated on held-out test sets. The clinTALL architecture was evaluated using identical splits, default TabM parameters, inverse class-frequency weighting, PLE-transformed numerical features, and AdamW optimization with early stopping.

Performance metrics were aggregated across all folds. Overall accuracy and a cumulative confusion matrix were computed to identify systematic misclassification patterns.

#### External Validation

For external validation, the clinTALL model was trained using the set of expressed genes shared between the training and validation cohorts, corresponding to the final gene set defined above. Hyperparameter tuning was performed exclusively on the training data. Raw RNA-seq counts were normalized using variance-stabilizing transformation (VST) implemented in pyDESeq2 [34], with normalization parameters estimated exclusively from the training cohort and applied to validation cohorts without refitting.

Five models trained during cross-validation were used to generate predictions for external cohorts, and final predictions were obtained by averaging outputs across models.

## Results

### Input of multimodal data achieves high performance of T-ALL subtype classification and treatment outcome inference

clinTALL is a multitask learning pipeline that integrates multimodal input data for pediatric T-ALL subtype classification and treatment outcome prediction (Fig. 1A). The model was trained on 1309 T-ALL samples from a recently published well-curated cohort of pediatric T-ALL patients [8]. Multimodal inputs contained a total of 2108 features collected at diagnosis, including 41 baseline clinical parameters and day-29 minimal residual disease (MRD), 1066 whole genome sequencing (WGS) derived genomic alterations and 1000 gene expression features from whole transcriptome sequencing (WTS, Supplementary Table S3). To process this heterogeneous tabular data, we utilized TabM ([18]), a modern deep learning architecture that employs piecewise linear embeddings and batch ensembling to learn robust feature representations. A neuronal network (NN) based architecture was employed to generate a shared latent embedding for jointly learned dual prediction of subtypes and risk scores (Fig. 1B). A softmax classification head was used for subtype prediction optimized via cross-entropy loss, whereas a cause-specific Cox Proportional Hazards (adapted from the DeepSurv implementation of the pycox package [35, 36]) approach was employed to generate risk scores, explicitly modeling competing risks (e.g., relapse vs. death) within a unified framework.

**Fig. 1.**
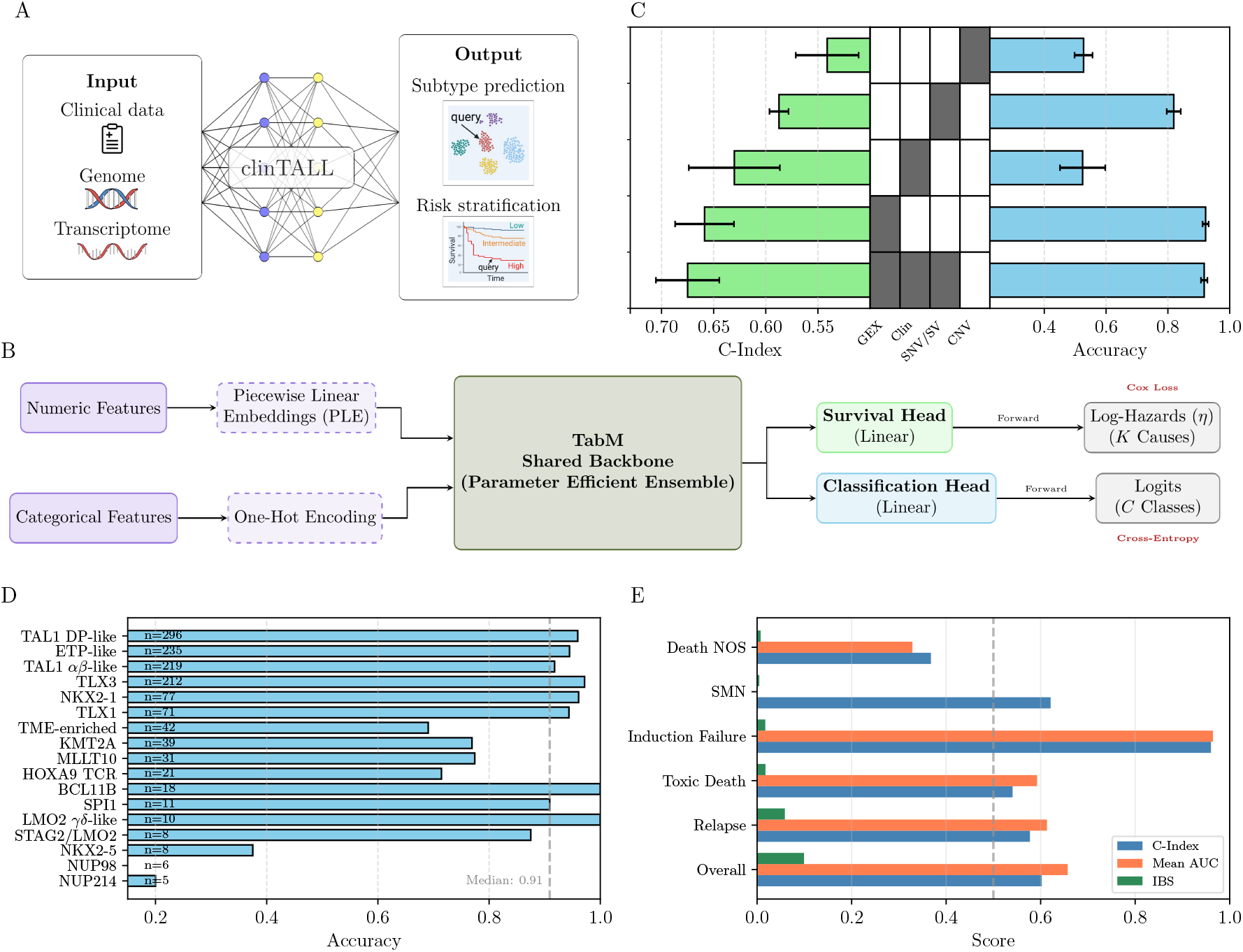
Overview and performance of the clinTALL multimodal framework. **A** Overview of the clinTALL framework. The model integrates clinical, genomic, transcriptomic, and CNV data and predicts both T-ALL molecular subtype and event-specific risk scores. **B** Schematic overview of the model architecture. **C** Performance comparison across different data modalities indicates that a multimodal data model outperforms unimodal data models. The central heatmap defines the input data configuration for each experiment, where cells colored in grey indicate the inclusion of a specific data modalities (Copy number variation (CNV), clinical data, genomic alterations (SNV/SV), or gene expression). The left bar chart displays the mean concordance index (C-Index) for survival prediction, and the right bar chart shows the mean accuracy for subtype classification. Error bars represent the standard deviation. **D** The per-subtype classification accuracy shows high predictive performance for most subtypes. Horizontal bars show recall (per-class accuracy) for each T-ALL subtype. The number of evaluation samples for each subtype is listed to the left. Most subtypes achieve high accuracy, whereas a small number of rare subtypes with very low sample counts show limited discriminability. **E** Performance comparison of three risk prediction metrics across different clinical event types. The C-Index (blue bars) measures discrimination ability, mean time-dependent AUC (orange bars) represents the average area under the receiver operating characteristic curve, and IBS (green bars, Integrated Brier Score) assesses prediction error, where lower values indicate better performance. A dashed vertical line at 0.5 represents random prediction for AUC-based metrics.

To evaluate the contribution of each data modality, we benchmarked single-modality data models. Among these, the transcriptome-only model showed the strongest predictive performance for both subtype classification (92.2%) and event-free survival (EFS) prediction (65.9%) (Fig. 1C, Supplementary Table S5). The high subtype-classification accuracy is expected, given that these subtypes were originally defined using gene expression based clustering [8]. WGS derived alterations also achieved good performance for both tasks. Clinical parameters demonstrated relatively low subtype classification accuracy (52.4%), however displayed good concordance for EFS prediction (63.0%). Together, these data indicate that the transcriptome-only model outperforms models solely based on genomic or clinical features across both tasks.

Integration of all data modalities preserves the high accuracy of subtype classification, and also achieves high accuracy at the individual subtype level (Fig. 1D). Among the 17 T-ALL subtypes, nine achieved an accuracy greater than 90%. Four subtypes (TME-enriched, KMT2A, MLLT10 and HOXA9 TCR) showed moderate accuracy (*<* 75%) due to overlapping features among these subtypes, while the TME-enriched subtype was further affected by reduced sample quality, with a lower blast percentage compared to the overall cohort (51.2% vs. 84.2%). Three subtypes (NKX2-5, NUP98 and NUP214) exhibited very low accuracy (*<* 40 %), which is likely due to the limited number of samples for training.

Instead of classical single-event based survival analysis, clinTALL employs a competing risk-based framework to generate event-specific risk predictions. The cause-specific competing risk model achieved an overall concordance-index (C-index) of 0.602, and demonstrated distinct predictive capabilities across event types (Fig. 1E, Table 1). Performance was the strongest for induction failure, where the model maintained high discrimination (C-index 0.960; mean AUC 0.965). Secondary outcomes, including relapse and second malignant neoplasm, showed moderate discriminative ability (C-indices of 0.577 and 0.621, respectively) accompanied by low Brier scores, indicating low overall prediction error despite the low event frequency.

**Table 1.**
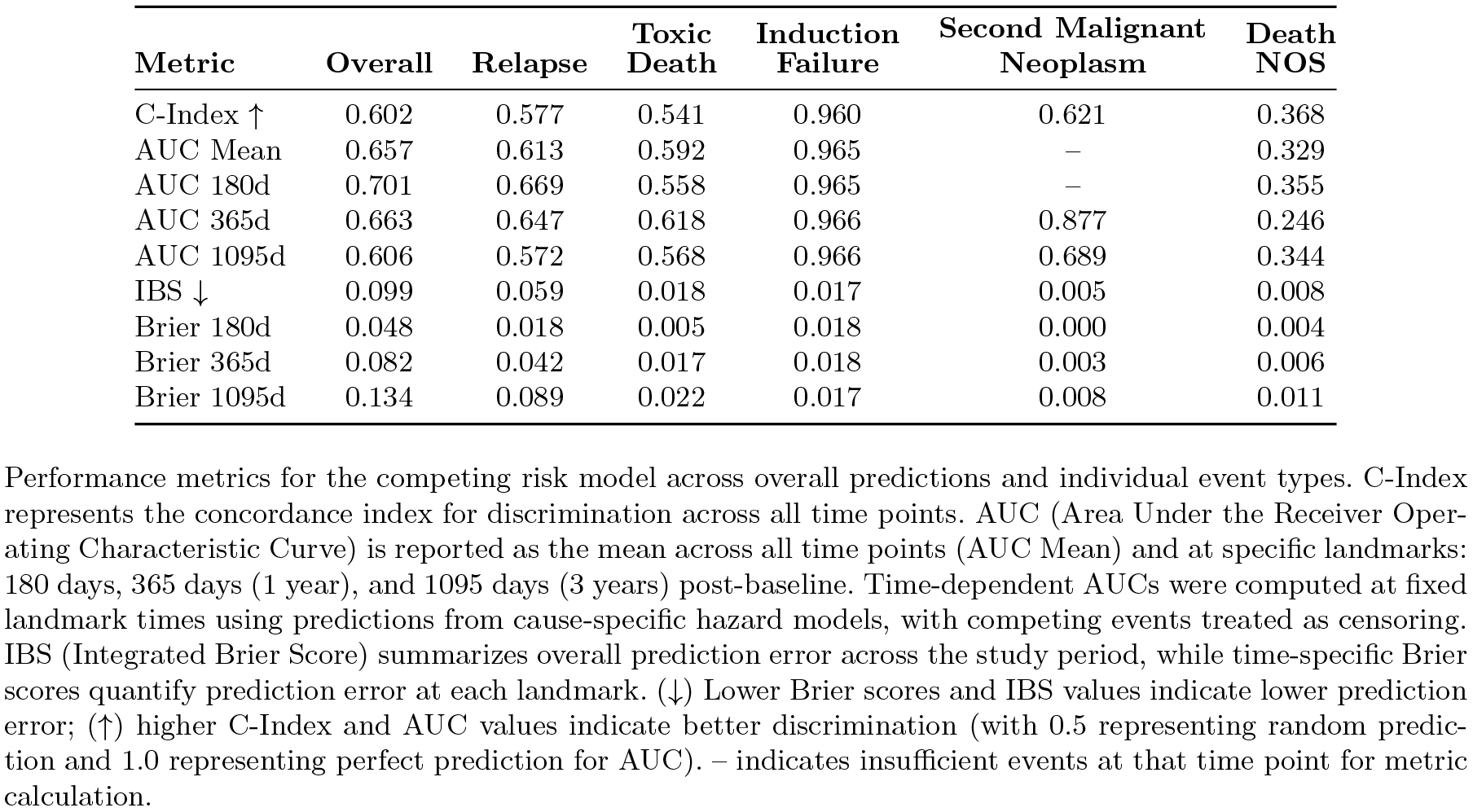
Comprehensive Performance Metrics of Competing Risk Models by Event Type.

| Metric | Overall | Relapse | Toxic Death | Induction Failure | Second Malignant Neoplasm | Death NOS |
| --- | --- | --- | --- | --- | --- | --- |
| C-Index ↑ | 0.602 | 0.577 | 0.541 | 0.960 | 0.621 | 0.368 |
| AUC Mean | 0.657 | 0.613 | 0.592 | 0.965 | – | 0.329 |
| AUC 180d | 0.701 | 0.669 | 0.558 | 0.965 | – | 0.355 |
| AUC 365d | 0.663 | 0.647 | 0.618 | 0.966 | 0.877 | 0.246 |
| AUC 1095d | 0.606 | 0.572 | 0.568 | 0.966 | 0.689 | 0.344 |
| IBS ↓ | 0.099 | 0.059 | 0.018 | 0.017 | 0.005 | 0.008 |
| Brier 180d | 0.048 | 0.018 | 0.005 | 0.018 | 0.000 | 0.004 |
| Brier 365d | 0.082 | 0.042 | 0.017 | 0.018 | 0.003 | 0.006 |
| Brier 1095d | 0.134 | 0.089 | 0.022 | 0.017 | 0.008 | 0.011 |

Performance metrics for the competing risk model across overall predictions and individual event types. C-Index represents the concordance index for discrimination across all time points. AUC (Area Under the Receiver Operating Characteristic Curve) is reported as the mean across all time points (AUC Mean) and at specific landmarks: 180 days, 365 days (1 year), and 1095 days (3 years) post-baseline. Time-dependent AUCs were computed at fixed landmark times using predictions from cause-specific hazard models, with competing events treated as censoring. IBS (Integrated Brier Score) summarizes overall prediction error across the study period, while time-specific Brier scores quantify prediction error at each landmark. ( ↓ ) Lower Brier scores and IBS values indicate lower prediction error; ( ↑ ) higher C-Index and AUC values indicate better discrimination (with 0.5 representing random prediction and 1.0 representing perfect prediction for AUC). – indicates insufficient events at that time point for metric calculation.

Together, by integrating clinical features and multi-omics data, clinTALL enables high performance subtype classification, and allows for event-specific risk prediction.

### Competing risk-based modeling determined patients at high risk for individual events

Pediatric T-ALL outcome data contain a set of mutually competing events, such as induction failure, toxic death and relapse. Thus, a competing risk-based survival analysis is required for accurate analysis of treatment outcome. Accordingly, clinTALL models survival of pediatric T-ALL using a competing risk structure and generates event specific risk scores. Using cumulative incidence functions we confirmed the model’s ability to distinguish high-risk patient groups across key clinical endpoints (Fig. 2A). The most pronounced risk stratification was observed for induction failure, where the model achieved near-perfect separation, isolating virtually all events within the predicted high-risk group. Second malignant neoplasm events showed moderate separation consistent with their lower baseline frequency, with the high-risk group consistently tracking above the low-risk group. Conversely, stratification for death not otherwise specified (death NOS) was unsuccessful (no patients with events ended up in the high-risk group), likely reflecting the heterogeneous nature of this small catch-all category.

**Fig. 2.**
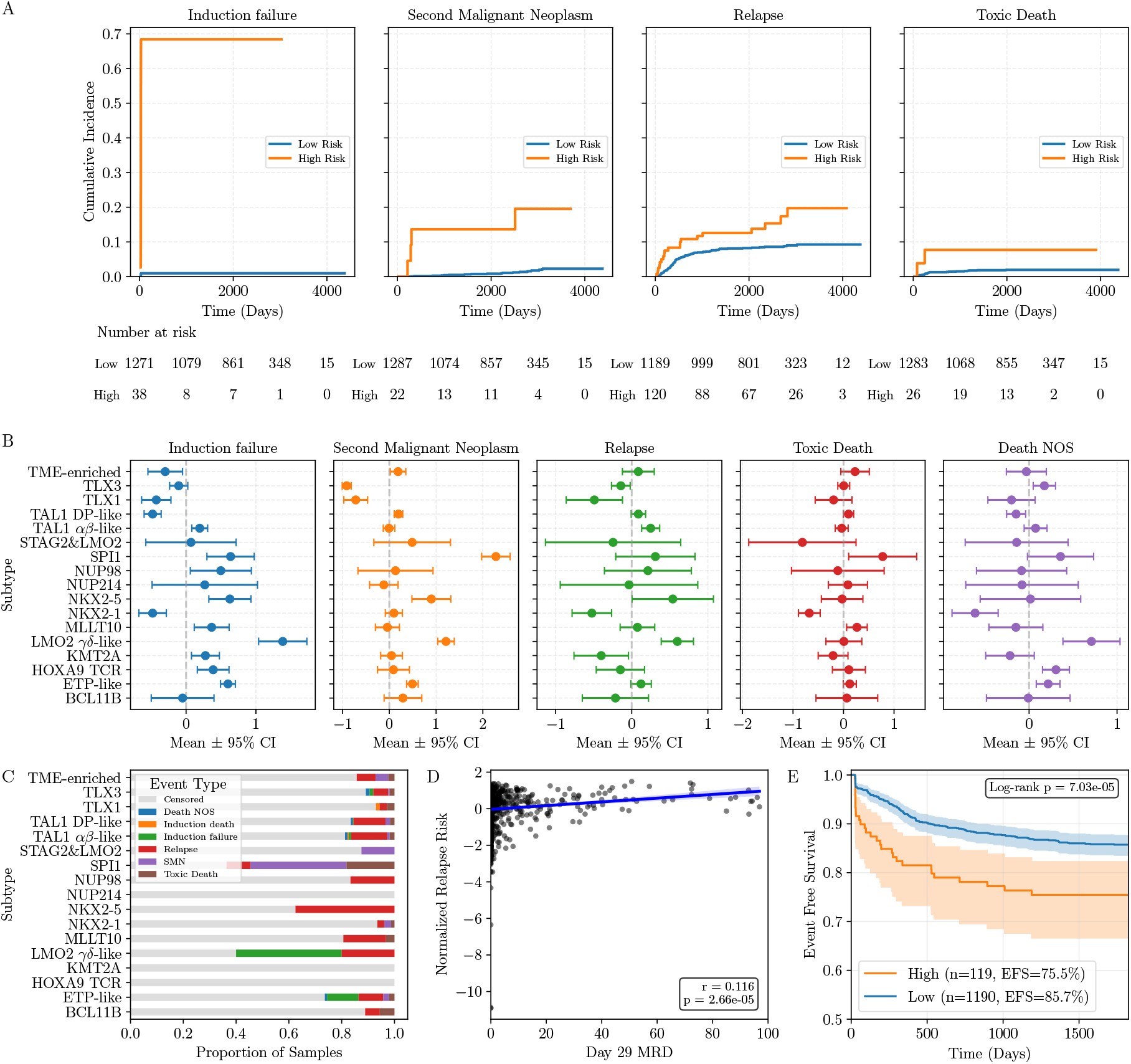
Integrated evaluation of competing risk model predictions. **A** Cumulative incidence curve stratified by predicted risk groups (high risk in orange, low risk in blue) for five competing event types. Patients were classified into risk groups based on model outputs at baseline. The y-axis represents the cumulative probability of experiencing each specific event over time (x-axis, in days from baseline), accounting for competing risks. The stepwise pattern reflects discrete event times in the dataset. Patients were classified as high-risk if their predicted cause-specific log-hazard was at or above the percentile threshold corresponding to the observed event rate for each specific event type. **B** Standardized cause-specific log-hazard coefficients with 95% confidence intervals from the competing risk model. Each panel displays coefficients scaled within that event type. **C** Proportion of samples experiencing each adverse event type, stratified by T-ALL subtype. Event types include: no event (grey), death not otherwise specified (NOS), induction death, induction failure, relapse, second malignant neoplasm, and toxic death. **D** Correlation between Day 29 minimal residual disease (MRD) levels (%) and normalized predicted relapse risk scores. Each point represents an individual patient sample. (r=0.116, p=2.66× 10^−5^, Pearson, two-tailed significance test) **E** 5-year event-free survival curves by predicted relapse-specific risk group. The high-risk group (orange, n=119) shows significantly inferior EFS compared to the low-risk group (blue, n=1,190). Shaded regions indicate 95% confidence intervals. Relapse-specific risk groups were defined using model-predicted probabilities with cutoffs matching observed event rates.

When applied to a multimodal dataset, our event-specific model enables a more refined characterization of patient outcomes and improved discrimination among samples experiencing different clinical endpoints. For example, previous studies using simple event-censor-based survival models have suggested that SPI1 and LMO *γδ*-like subtypes are associated with poor treatment outcome [8, 37]. In our competing risk-based model, we observed that cases with LMO2 *γδ*-like subtype show specific increased risk scores for induction failure and relapse, whereas cases with SPI1 subtype were associated with second malignancy and toxic death (Fig. 2B). This is in line with the enriched frequency of these specific clinical events in cases harbouring LMO2 *γδ*-like and SPI1 T-ALL subtypes (Fig. 2C). Together, this highlights the superior accuracy and resolution of clinTALL in event-specific outcome inference.

Outcomes for relapsed pediatric T-ALL remained poor [37], therefore identification of patients at high risk for relapse is critical to enable precise therapeutic stratification. At present, minimal residual disease (MRD) remains the only prognostic marker applied across pediatric T-ALL clinical trials [38, 39]. In our competing risk-based model, although C-index (57.7%) for relapse is modest relative to other events, it demonstrates strong short-term discrimination (180-day AUC = 0.669), with attenuation over longer horizons (1-year AUC = 0.647, 3-year AUC = 0.572, Table 1). The relapse risk score derived from our model is correlated positively with day-29 MRD (r = 0.116, p = 2.66×10^−5^; Fig. 2D, Table 1). Moreover, our competing risk-based model successfully delineated two distinct risk groups, with the high-risk group exhibiting a significantly steeper accumulation of early events and a higher cumulative incidence plateau (∼20% vs. ∼10%; Fig. 2A). Consistently, high-risk patients showed significantly lower 5-years EFS (75.5% vs. 85.7%, p = 7.03× 10^−5^; Fig. 2E). Together, these findings indicate that the competing risk-based model is able to identify patients at high-risk to relapse.

### clinTALL outperforms previous tool in subtype prediction

We next compared the performance of clinTALL with previously published tools. To our knowledge, no existing model provides both subtype classification and outcome prediction for pediatric T-ALL. For outcome prediction alone, PredicTALL has been reported [15], but its source code is not publicly available, preventing direct comparison with clinTALL. Several AI-based tools have been published for ALL subtype prediction, though most address pan-ALL subtype classification and do not specifically resolve the complex heterogeneity of T-ALL [10, 14, 40]. By contrast, TALLsorts and Tallforest were reported to capture the complex subtype architecture of T-ALL [9, 13]. Among these, only TALLsorts provides accessible source code; we therefore compared clinTALL with TALLsorts for subtype prediction performance. To enable a direct comparison with TALLsorts that only uses gene expression data as input, we retrained its model using the same cohort of 1309 T-ALL samples and the same 17 subtype output as applied in clinTALL. Under these harmonized conditions, clinTALL outperformed TALLsorts, with an overall accuracy of 95.0 % versus 93.4 % (Fig. 3A). Furthermore, clinTALL showed higher per-class accuracy (macro recall of 86.1 % vs. 83.4 %; Fig. 3B), with notably improved performance for LMO2 *γδ*-like (90.0 % vs. 82.0 %) and MLLT10 (90.0 % vs. 86.0 %) subtypes (Fig. 3C-D). Together, despite the fact that only RNA-sequencing data were used for training, clinTALL outperforms other available tools for subtype prediction.

**Fig. 3.**
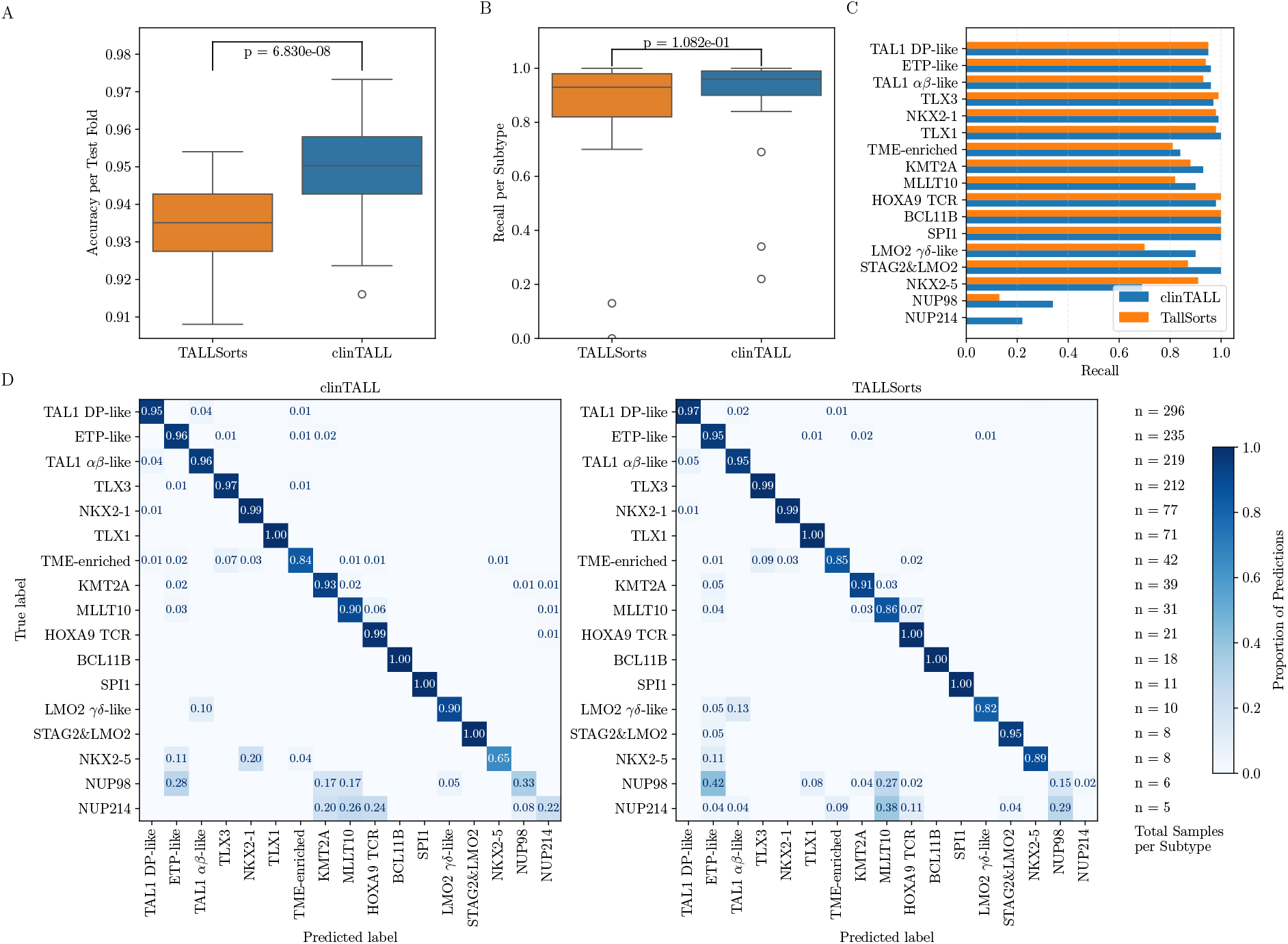
clinTALL demonstrates superior accuracy in T-ALL subtype classification compared to TallSorts. **A** Overall classification accuracy per test fold. clinTALL achieves significantly higher accuracy than TallSorts (p-value:6.83× 10^−8^, paired t-test). Box plots show median and interquartile ranges across test folds. **B** Overall recall (sensitivity) comparison between clinTALL and TallSorts. No significant difference in overall recall was observed (p-value: 0.102, paired t-test). **C** Subtype-specific recall comparison. Bar plots show the proportion of correctly classified samples (recall) for each of the 17 T-ALL molecular subtypes. clinTALL (orange) demonstrates improved or comparable performance to TallSorts (blue) across most subtypes, with notable improvements in rare subtypes. **D** Confusion matrices showing classification performance for both methods. Left panel: clinTALL predictions. Right panel: TallSorts predictions. Rows represent true subtype labels, columns represent predicted labels. Values indicate the proportion of samples assigned to each predicted category. Sample sizes for each subtype are shown on the right (n=5 to n=296).

### High accuracy was observed in external cohorts

To estimate the performance of clinTALL on real-world clinical data, we applied the tool to two independent RNA-sequencing datasets from diagnostic pediatric T-ALL samples. The Hackenhaar dataset comprises 108 T-ALL samples from the Nordic Society of Paediatric Haematology (NOPHO) ALL2008 T-ALL study cohort [22]. However, the absence of publicly available clinical or cytogenetic annotations prevented the direct assessment of prediction accuracy. The second, in-house RNA-seq dataset consists of 120 diagnostic samples from patients with T-ALL enrolled in the Italian AIEOP-BFM ALL 2017 treatment study. In this cohort, the exact fusion gene was assessed positive for 41 samples, whereas the others were negative. Given the fact that the 17 subtypes defined in the Pölönen study [8] are derived from integrated whole-transcriptome and whole-genome analyses, the lack of definitive ground truth subtype information in these two independent RNA-seq datasets closely mirrors the incompleteness and heterogeneity of real-world diagnostic data.

First, despite missing ground truth labels, we sought to estimate whether subtypes predicted by clinTALL aligned with the structure of the training cohort. To this end, we embedded both the query and reference samples into a shared UMAP space. As a result, we observed strong concordance between the predicted subtypes and their corresponding reference regions (Fig. 4A), indicating that clinTALL maintains robust performance when applied to real-world data.

**Fig. 4.**
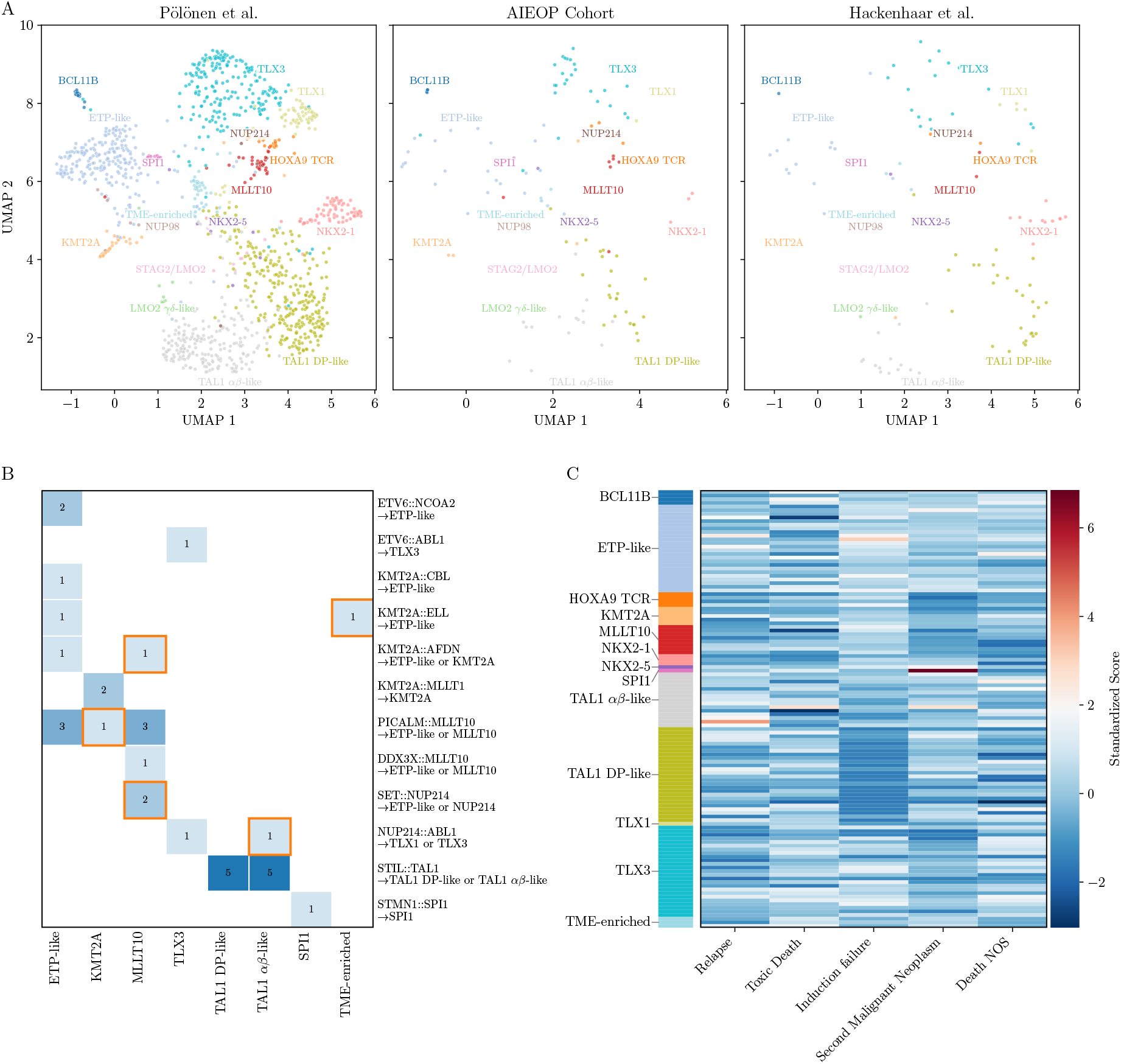
External validation of clinTALL subtype classification and risk prediction across independent cohorts. **A** UMAP projection showing the distribution of T-ALL molecular subtypes across three independent cohorts: Pölonen et al. (left), cohort from AIEOP-BFM ALL 2017 study, and Hackenhaar et al. (right). Each point represents an individual patient sample, colored by predicted subtype. The cohorts demonstrate consistent subtype distributions in transcriptomic space, validating the generalizability of clinTALL classification. **B** Validation of clinTALL subtype predictions for samples with known fusion events. The confusion matrix shows the number of samples with specific gene fusions (rows) assigned to each predicted subtype (columns). Results demonstrate biologically concordant classifications, with fusion-driven cases correctly assigned to their expected subtypes (e.g., *STIL*::*TAL1* → TAL1 DP-like or TAL1 *αβ*-like; *KMT2A* fusions → KMT2A or ETP-like subtypes; *PICALM* ::*MLLT10* and *DDX3X* ::*MLLT10* → MLLT10 or ETP-like). **C** Visualisation of the standardized estimated log-hazard for different cause-specific adverse clinical endpoints (X-axis) across individual patients (Y-axis) in the in-house validation cohort. Each row represents a single patient, ordered and grouped by their predicted T-ALL molecular subtype (indicated by the colored bar on the left).

Next, we evaluated the prediction accuracy using the available fusion gene information in the AIEOP validation cohort. Among the 20 fusion gene types determined in the in-house cohort, twelve were also reported in the Pölönen study, allowing us to manually assign subtypes according to their definition in 33 samples (Supplementary Table S6-7). Of these samples, clinTALL correctly predicted subtypes in 27 samples, corresponding to an accuracy of 81.8 % (Fig. 4B). The false predicted samples are mainly from the NUP214 and related subtypes, mainly due to the low number of samples in the training data (Fig. 4B).

Lastly, we assessed the accuracy of predicting treatment outcomes in the Italian validation cohort with only RNA-sequencing data available. We determined two patients with a high risk score for second malignant neoplasm (one SPI1 and one TAL1ab-like) and one ETP-like patient with a high risk score of induction failure (Fig. 4C). In addition, five patients were classified as high-risk to relapse (Fig. 4C). Following treatment outcome assessment, these predicted outcomes were confirmed in all eight patients.

These findings demonstrate that clinTALL accurately predicts molecular T-ALL subtypes using real-world RNA-sequencing data and allows for determination of patients with high risk for induction failure, second malignancy and relapse.

### Container-based implementation for clinical deployment

To enable clinical translation of clinTALL, we provide an interactive Docker-based application for subtype and risk prediction. The application accepts standard clinical and/or molecular features and generates patient-specific predictions for each competing event. The tool is designed for ease of use by clinicians without programming expertise and supports both single-patient queries and batch processing. Comprehensive documentation and example cases are provided to facilitate interpretation of model outputs. The clinTALL application is freely accessible on GitHub.

## Discussion

Despite the increasingly refined molecular classification of pediatric T-ALL, translating this knowledge into clinically applicable tools remains challenging. In this study, we developed clinTALL, a ML-based framework that integrates multi-modal data for both genetic subtype classification and treatment outcome prediction. By incorporating clinical, genomic, and transcriptomic features, clinTALL consistently outperformed single-modality models in both tasks. Notably, while we observed that transcriptomic features alone were central to capturing T-ALL biology, the integration of multi-omics data via the TabM architecture successfully leveraged complementary information without introducing noise. This approach preserved the high fidelity of RNA-seq-based subtype classification while simultaneously enhancing survival prediction capabilities.

The competing risk-based modeling provided a particularly effective approach for stratifying distinct clinical endpoints, though performance varied by event type. clinTALL was highly effective at identifying induction failure and second malignant neoplasm, yet performance was lower for clinically heterogeneous categories such as toxic death. Regarding relapse, although clinTALL achieved strong concordance for early events, performance gradually declined at later time points. This trend likely reflects the temporal differences in these events: induction failure is defined at the end of the 4-6 week induction phase [41–43], whereas relapse events occur at a median of 13.8 months [44]. Nevertheless, the biological validity of the model was supported by an observed correlation between predicted relapse risk and day-29 MRD. This suggests that clinTALL captures intrinsic components of early treatment response. While clinTALL preserves its predictive power for a disease typically associated to early events, however, the reduced accuracy at later time points indicates that early features alone may be insufficient to capture the biological process evolving during treatment, or processes that may drive late relapses, suggesting that incorporating longitudinal molecular or clinical measurements may be required to improve long-term predictive accuracy. Although clinTALL achieved high subtype accuracy overall, four subtypes exhibited reduced performance due to the limited number of samples in the training cohort. Expanding the sample numbers for these rare subtypes will help to further refine the performance. Similarly, event-specific risk modeling would also benefit from validation in datasets with complete and accurately annotated time of clinical events.

## Conclusion

In summary, clinTALL provides a unified multimodal, multitask framework that simultaneously delivers subtype classification and event-specific risk prediction in pediatric T-ALL. By offering a user-friendly web interface, clinTALL facilitates the broader application of multi-omics-derived knowledge in clinical research. This framework supports the integration of molecular information into patient management, ultimately contributing to the development of more tailored treatment options and improved treatment outcomes.

## Abbreviations

T-LL: T-lineage acute lymphoblastic leukemia
B-ALL: B-lineage acute lymphoblastic leukemia
ETP-ALL: Early T-cell precursor acute lymphoblastic leukemia
WHO: World Health Organization
ICC: International Consensus Classification
ML: Machine learning
EFS: Event-free survival
CNV: Copy number variation
SNV: Single nucleotide variant
SV: Structural variant
C-index: Concordance index
AUC: Area under the curve
IBS: Integrated Brier score
WGS: Whole-genome sequencing
WTS: Whole-transcriptome sequencing
NN: Neural network
PLE: Piecewise linear embeddings
NOS: Not otherwise specified
MRD: Minimal residual disease
CIF: Cumulative incidence function

## Supplementary information

Additional File ‘Supplementary.xlsx’ includes:

Table S1 - Genes selected in supervised selection process

Table S2 - Number of features used per model

Table S3 - Names of features per model

Table S4 - Final model parameters

Table S5 - Final model classification reports

Table S6 - Fusion gene to subtype assignement for validation

Table S7 - Predictions and fusion genes for validation cohort

## Acknowledgements

We would like to thank Dr. Matt McCrary for critical reading of the manuscript and insightful comments.

## Declarations

### Author’s Contributions

Conceptualization: ŽA, JY and AKB. Methodology: LS and JY. Pipeline development: LS and JY. Validation: ŽA, SR, GF, AB, FL, CR and GiC. Data curation: SR, GF, AR, LL, GuC, AB, FL, CR, GiC. Writing - Original draft preparation: LS, JY. Writing-Review and editing: LS, ŽA, JY and AKB. Supervision: AKB. Project administration: AKB. Funding acquisition: GiC and AKB. All authors read and approved the final manuscript.

### Data availability

The training datasets used during the current study are available from Synapse (ID: syn54032669; https://doi.org/10.7303/syn54032669) and the study by Pölönen et al. [8]. The NOPHO ALL2008 validation dataset from the study by Hackenhaar et al. [22] is available from Gene Expression Omnibus (GEO) under accession number GSE272023. The RNA-seq derived count matrix for the AIEOP cohort can be provided upon request.

### Material availability

Not applicable

### Code availability

The code accompanying this paper is available at https://github.com/UKWgenommedizin/clinTALL/ and Zenodo

### Competing interests

The authors declare that they have no competing interests.

### Consent for publication

Not applicable

### Ethics approval and consent to participate

Not applicable

### Funding

This project was supported by the BZKF as part of the OMICS and AI/Bioinformatics flagship programs. This research was partly funded by Deutsche Forschungsgemeinschaft (DFG), grant numbers BE6555/1-1 and BE6555/2-1. This work was supported by a grant of the Mildred Scheel Early Career Center (MSNZ Würzburg) to Željko Antić. This project was also supported by Fondazione Maria Letizia verga.

## Notes

### Competing Interest Statement

The authors have declared no competing interest.

## References

[1] Shiraz P, Jehangir W, Agrawal V. T-Cell Acute Lymphoblastic Leukemia-Current Concepts in Molecular Biology and Management. Biomedicines. 2021 11;9. 10.3390/biomedicines9111621.

[2] Summers RJ, Teachey DT. SOHO State of the Art Updates and Next Questions — Novel Approaches to Pediatric T-cell ALL and T-Lymphoblastic Lymphoma. Clinical lymphoma, myeloma & leukemia. 2022 10;22:718–725. 10.1016/j.clml.2022.07.010.

[3] Brady SW, Roberts KG, Gu Z, Shi L, Pounds S, Pei D, et al. The genomic landscape of pediatric acute lymphoblastic leukemia. Nature genetics. 2022 9;54:1376–1389. 10.1038/s41588-022-01159-z.

[4] Arber DA, Orazi A, Hasserjian RP, Borowitz MJ, Calvo KR, Kvasnicka HM, et al. International Consensus Classification of Myeloid Neoplasms and Acute Leukemias: integrating morphologic, clinical, and genomic data. Blood. 2022 9;140:1200–1228. 10.1182/blood.2022015850.

[5] Liu Y, Easton J, Shao Y, Maciaszek J, Wang Z, Wilkinson MR, et al. The genomic landscape of pediatric and young adult T-lineage acute lymphoblastic leukemia. Nature genetics. 2017 8;49:1211–1218. 10.1038/ng.3909.

[6] Dai YT, Zhang F, Fang H, Li JF, Lu G, Jiang L, et al. Transcriptome-wide subtyping of pediatric and adult T cell acute lymphoblastic leukemia in an international study of 707 cases. Proceedings of the National Academy of Sciences of the United States of America. 2022 4;119:e2120787119. 10.1073/pnas.2120787119.

[7] Mü ller J, Walter W, Haferlach C, Mü ller H, Fuhrmann I, Müller ML, et al. How T-lymphoblastic leukemia can be classified based on genetics using standard diagnostic techniques enhanced by whole genome sequencing. Leukemia. 2023 1;37:217–221. 10.1038/s41375-022-01743-6.

[8] Pölönen P, Giacomo DD, Seffernick AE, Elsayed A, Kimura S, Benini F, et al. The genomic basis of childhood T-lineage acute lymphoblastic leukaemia. Nature. 2024 8;632:1082–1091. 10.1038/s41586-024-07807-0.

[9] Gu A, Schmidt B, Lonsdale A, Jalaldeen R, Kosasih HJ, Brown LM, et al. TALL-Sorts: a T-cell acute lymphoblastic leukemia subtype classifier using RNA-seq expression data. Blood advances. 2023 12;7:7402–7406. 10.1182/bloodadvances.2023010385.

[10] Krali O, Marincevic-Zuniga Y, Arvidsson G, Enblad AP, Lundmark A, Sayyab S, et al. Multimodal classification of molecular subtypes in pediatric acute lymphoblastic leukemia. NPJ precision oncology. 2023 12;7:131. 10.1038/s41698-023-00479-5.

[11] Tang M, Antić Ž, Fardzadeh P, Pietzsch S, Schröder C, Eberhardt A, et al. An artificial intelligence-assisted clinical framework to facilitate diagnostics and translational discovery in hematologic neoplasia. EBioMedicine. 2024 6;104:105171. 10.1016/j.ebiom.2024.105171.

[12] Gu Z, Hu Z, Jia Z, Liu J, Mao A, Han H. MD-ALL: an Integrative Platform for Molecular Diagnosis of B-cell Acute Lymphoblastic Leukemia. Research square. 2023 4;10.21203/rs.3.rs-2798895/v1.

[13] Polonen P, Fan Y, Lei S, Gao Q, Wu Y, Chang TC, et al. Tallforest: Multi-omic classifier for T-lineage acute lymphoblastic leukemia. Blood. 2025 11;146:336–336. 10.1182/blood-2025-336.

[14] Steinicke TL, Benfatto S, Capilla-Guerra MR, Monteleone AB, Young JH, Shankar S, et al. Rapid epigenomic classification of acute leukemia. Nature genetics. 2025 10;57:2456–2467. 10.1038/s41588-025-02321-z.

[15] Simonin M, Boissel N, Petit A, Mullighan CG, Hunger SP, Loh ML, et al. Prognostic impact of the PredicT-ALL classifier in the AALL0434 trial: a model combining NGS, MRD, and WBC at diagnosis. Blood advances. 2025 10;9:5323–5326. 10.1182/bloodadvances.2025017399.

[16] Satagopan JM, Ben-Porat L, Berwick M, Robson M, Kutler D, Auerbach AD. A note on competing risks in survival data analysis. British journal of cancer. 2004 10;91:1229–35. 10.1038/sj.bjc.6602102.

[17] Buckley M, Yeung DT, White DL, Eadie LN. T-cell acute lymphoblastic leukaemia: subtype prevalence, clinical outcome, and emerging targeted treatments. Leukemia. 2025 6;39:1294–1310. 10.1038/s41375-025-02599-2.

[18] Gorishniy Y, Kotelnikov A, Babenko A. TabM: Advancing Tabular Deep Learning with Parameter-Efficient Ensembling. arXiv preprint arXiv:241024210. 2025;10.48550/arXiv.2410.24210.

[19] Andrews S.: FastQC: A Quality Control Tool for High Throughput Sequence Data. https://www.bioinformatics.babraham.ac.uk/projects/fastqc/.

[20] Dobin A, Davis CA, Schlesinger F, Drenkow J, Zaleski C, Jha S, et al. STAR: ultrafast universal RNA-seq aligner. Bioinformatics. 2012 10;29(1):15–21. 10.1093/bioinformatics/bts635.

[21] Liao Y, Smyth GK, Shi W. featureCounts: an efficient general purpose program for assigning sequence reads to genomic features. Bioinformatics. 2013 11;30(7):923–930. 10.1093/bioinformatics/btt656.

[22] Hackenhaar FS, Refhagen N, Hagleitner M, van Leeuwen F, Marquart HV, Madsen HO, et al. CpG island methylator phenotype classification improves risk assessment in pediatric T-cell acute lymphoblastic leukemia. Blood. 2025 5;145:2161–2178. 10.1182/blood.2024026027.

[23] Pedregosa F, Varoquaux G, Gramfort A, Michel V, Thirion B, Grisel O, et al. Scikitlearn: Machine Learning in Python. Journal of Machine Learning Research. 2011;12:2825– 2830.

[24] Lundberg SM, Erion G, Chen H, DeGrave A, Prutkin JM, Nair B, et al. From local explanations to global understanding with explainable AI for trees. Nature Machine Intelligence. 2020;2(1):2522–5839.

[25] Gorishniy Y, Rubachev I, Babenko A. On Embeddings for Numerical Features in Tabular Deep Learning. arXiv preprint arXiv:220305556. 2023;10.48550/arXiv.2203.05556.

[26] Loshchilov I, Hutter F. Decoupled Weight Decay Regularization. In: International Conference on Learning Representations; 2019. Paper ID: Bkg6RiCqY7. Available from: https://openreview.net/forum?id=Bkg6RiCqY7.

[27] Akiba T, Sano S, Yanase T, Ohta T, Koyama M. Optuna: A Next-generation Hyperparameter Optimization Framework. In: Proceedings of the 25th ACM SIGKDD International Conference on Knowledge Discovery and Data Mining; 2019. .

[28] Paszke A, Gross S, Chintala S, Chanan G, Yang E, DeVito Z, et al. Automatic differentiation in PyTorch. In: NIPS-W; 2017.

[29] Pölsterl S. scikit-survival: A Library for Time-to-Event Analysis Built on Top of scikit-learn. Journal of Machine Learning Research. 2020;21(212):1–6.

[30] Virtanen P, Gommers R, Oliphant TE, Haberland M, Reddy T, Cournapeau D, et al. SciPy 1.0: Fundamental Algorithms for Scientific Computing in Python. Nature Methods. 2020;17:261–272. s41592-019-0686-2.10.1038/

[31] Davidson-Pilon C. lifelines: survival analysis in Python. Journal of Open Source Software. 2019;4(40):1317. 10.21105/joss.01317.

[32] Hunter JD. Matplotlib: A 2D graphics environment. Computing in Science & Engineering. 2007;9(3):90–95. 10.1109/MCSE.2007.55.

[33] Waskom ML. seaborn: statistical data visualization. Journal of Open Source Software. 2021;6(60):3021. 10.21105/joss.03021.

[34] Muzellec B, Telenczuk M, Cabeli V, Andreux M. PyDESeq2: a python package for bulk RNA-seq differential expression analysis. Bioinformatics. 2023; 10.1093/bioinformatics/btad547.

[35] Katzman JL, Shaham U, Cloninger A, Bates J, Jiang T, Kluger Y. DeepSurv: personalized treatment recommender system using a Cox proportional hazards deep neural network. BMC Medical Research Methodology. 2018 12;18:24. 10.1186/s12874-018-0482-1.

[36] Kvamme H, ørnulf Borgan, Scheel I. Timeto-Event Prediction with Neural Networks and Cox Regression. Journal of Machine Learning Research. 2019;20(129):1–30.

[37] Hughes AD, Pölönen P, Teachey DT. Relapsed childhood T-cell acute lymphoblastic leukemia and lymphoblastic lymphoma. Haematologica. 2025 9;110:1934–1950. 10.3324/haematol.2024.285643.

[38] Wood BL, Devidas M, Summers RJ, Chen Z, Asselin B, Rabin KR, et al. Prognostic significance of ETP phenotype and minimal residual disease in T-ALL: a Children’s Oncology Group study. Blood. 2023 12;142:2069–2078. 10.1182/blood.2023020678.

[39] Cario G, Valsecchi MG, Conter V, Gotti G, Möricke A, Stanulla M, et al. Results in pediatric T-ALL patients treated in trial AIEOP-BFM ALL 2009: Prognostic factors in the context of modern risk-adapted therapy. HemaSphere. 2025;9(9):e70206. 10.1002/hem3.70206.

[40] Wiklander ML, Zachariah D, Krali O, Nordlund J. Error Reduction in Leukemia Machine Learning Classification With Conformal Prediction. JCO clinical cancer informatics. 2025 5;9:e2400324. 10.1200/CCI-24-00324.

[41] Inaba H, Greaves M, Mullighan CG. Acute lymphoblastic leukaemia. Lancet (London, England). 2013 6;381:1943–55. 10.1016/S0140-6736(12)62187-4.

[42] Stock W, Luger SM, Advani AS, Yin J, Harvey RC, Mullighan CG, et al. A pediatric regimen for older adolescents and young adults with acute lymphoblastic leukemia: results of CALGB 10403. Blood. 2019 4;133:1548–1559. 10.1182/blood-2018-10-881961.

[43] Maloney KW, Devidas M, Wang C, Mattano LA, Friedmann AM, Buckley P, et al. Outcome in Children With Standard-Risk B-Cell Acute Lymphoblastic Leukemia: Results of Children’s Oncology Group Trial AALL0331. Journal of clinical oncology : official journal of the American Society of Clinical Oncology. 2020 2;38:602–612. 10.1200/JCO.19.01086.

[44] Rheingold SR, Bhojwani D, Ji L, Xu X, Devidas M, Kairalla JA, et al. Determinants of survival after first relapse of acute lymphoblastic leukemia: a Children’s Oncology Group study. Leukemia. 2024 11;38:2382–2394. 10.1038/s41375-024-02395-4.

